# *Nf1* Deficiency Increases Mammary Collagen Deposition and Restricts Adipocyte Differentiation Before Tumor Formation

**DOI:** 10.1101/2023.06.01.539442

**Authors:** Menusha Arumugam, Elizabeth A. Tovar, Curt J. Essenburg, Patrick S. Dischinger, Ian Beddows, Emily Wolfrum, Zach B. Madaj, Lisa Turner, Kristin Feenstra, Kristin L. Gallik, Lorna Cohen, Madison Nichols, Rachel T.C. Sheridan, Corinne R. Esquibel, Ghassan Mouneimne, Carrie R. Graveel, Matthew R. Steensma

## Abstract

**BACKGROUND:** Neurofibromin, coded by the *NF1* tumor suppressor gene, is the main negative regulator of the RAS pathway and is frequently mutated in various cancers. Women with Neurofibromatosis Type I (NF1) – a tumor predisposition syndrome caused by a germline *NF1* mutation – have an increased risk of developing aggressive breast cancer with poorer prognosis. The mechanism by which *NF1* mutations lead to breast cancer tumorigenesis is not well understood. Therefore, the objective of this work was to identify stromal alterations before tumor formation that result in the increased risk and poorer outcome seen among NF1 patients with breast cancer.

**METHODS:** To accurately model the germline monoallelic *NF1* mutations in NF1 patients, we utilized an *Nf1-*deficient rat model with accelerated mammary development before presenting with highly penetrant breast cancer.

**RESULTS:** We identified increased collagen content in *Nf1*-deficient rat mammary glands before tumor formation that correlated with age of tumor onset. Additionally, gene expression analysis revealed that *Nf1*-deficient mature adipocytes in the rat mammary gland have increased collagen expression and shifted to a fibroblast and preadipocyte expression profile. This alteration in lineage commitment was also observed with *in vitro* differentiation, however, flow cytometry analysis did not show a change in mammary adipose-derived mesenchymal stem cell abundance.

**CONCLUSION:** Collectively, these studies uncovered the previously undescribed role of *Nf1* in mammary collagen deposition and regulating adipocyte differentiation. In addition to unraveling the mechanism of tumor formation, further investigation of adipocytes and collagen modifications in preneoplastic mammary glands will create a foundation for developing early detection strategies of breast cancer among NF1 patients.

**GRAPHICAL ABSTRACT:** 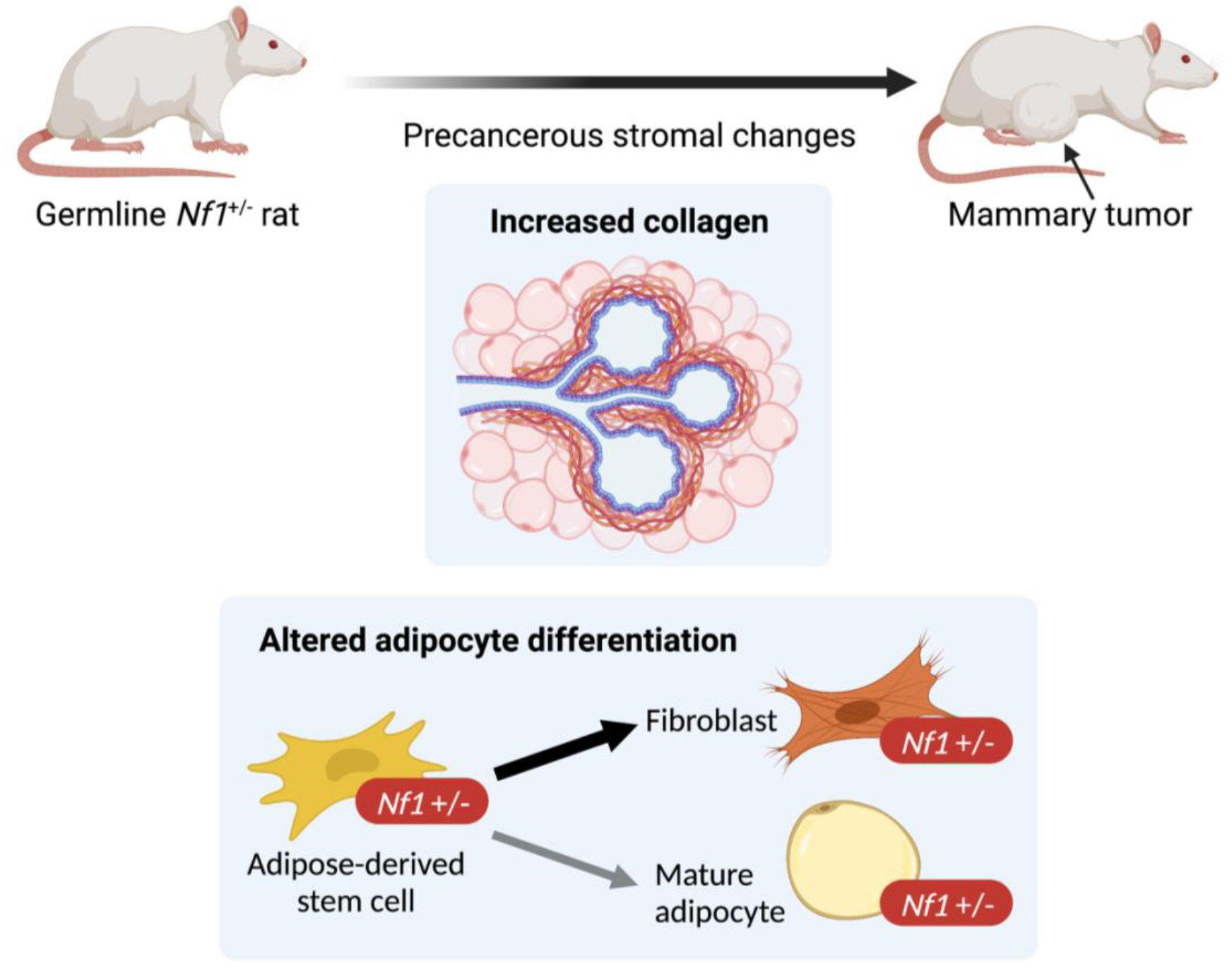

## BACKGROUND

Neurofibromatosis type 1 (NF1) is a tumor predisposition syndrome affecting 1 in 3,000 individuals worldwide with a wide range of clinical manifestations including cancer, neurological, cardiovascular, and musculoskeletal symptoms. (1-4) Cancer is the most common cause of death among NF1 individuals and reduces life expectancy by 10-15 years. This increased risk for developing cancer is due to mutation or deletion of the *Neurofibromin 1 (NF1)* gene, which disrupts neurofibromin tumor suppressor function as a key negative regulator of RAS. Neurofibromin, a 250-kDa protein, is the main negative regulator of RAS and prevents the activation of downstream RAS signaling, including the MAPK (mitogen-activated protein kinase) pathway. (5) Therefore, when *NF1* is mutated, and consequently unable to negatively regulate RAS, this results in increased transcription of genes involved in various cellular processes such as proliferation, differentiation, and apoptosis. (4, 6)

NF1 patients have a 5-7-fold increased risk of developing aggressive breast cancer with poorer prognosis compared to the general population. (7-12) This risk is also particularly higher at a younger age and has warranted the recommendation for earlier breast cancer screening at the age of 30 for women with NF1. (13, 14) Our analysis of the METABRIC breast cancer dataset showed that patients with at least one deleted copy of *NF1* are 1.65 times more likely to die within the first 10 years compared to patients with diploid *NF1* status. (15) Due to its critical tumor suppressor and driver mutation role in breast tumorigenesis, *NF1* is now considered a breast cancer susceptibility gene. (16) Despite this, the molecular mechanisms of *NF1*-mediated breast cancer formation are still poorly understood.

Although breast cancer arises from the epithelial cells, numerous studies have demonstrated the significant impact of the stroma on the earliest stages of tumor initiation, tumor progression, metastasis, and even clinical outcome. (17-20) Separated by a layer of basement membrane, the epithelial ducts are surrounded by stroma cells such as fibroblasts, adipocytes, endothelial cells, and immune cells embedded in a complex web of extracellular matrix (ECM). (21) In sporadic breast cancer, only epithelial cells harbor oncogenic mutations, but NF1 patients have germline mutations in their stromal and epithelial cells. Current studies on epithelial cells in *NF1*-related breast cancer have expanded our knowledge of their tumorigenic mechanisms, but there remains a gap in understanding of oncogenic contribution from an *NF1*-deficient stromal environment. The preneoplastic stromal alternations of inherited cancer syndromes have also been recently characterized in the context of individuals with germline BRCA1 mutations. This study identified a population of BRCA^+/mut^ fibroblast present before cancer formation that increased epithelial proliferation, altered epithelial differentiation, and promoted BRCA1-driven tumorigenesis *in vivo.* (22) Taking all this together, we do not know the extent of which *NF1*-deficient stromal cells provide a protumorigenic niche for epithelial transformation, thus contributing to the increased risk of breast cancer in individuals with NF1.

The current state of knowledge regarding the breast cancer stroma is heavily focused on cancer-associated fibroblasts (CAFs) and immune cells due to their potential as a therapeutic target. Despite the mammary gland being an adipocyte-dominant organ, we have not extensively studied the changes mammary adipocytes undergo during normal physiological breast remodeling and breast cancer. Contradicting the traditional view that mature adipocytes are terminally differentiated and non-proliferative, adipocytes can proliferate and differentiate into different cell types to adapt with the rapidly changing ductal epithelium structures during pregnancy, lactation, and involution. (23-26) Adipocytes can dedifferentiate to become a preadipocyte/mesenchymal-like cells through a process called adipocyte mesenchymal transition (AMT), a process shown to create a tumor-promoting environment. (27) In cancer, bidirectional crosstalk between tumor cells and stromal cells can push surrounding adipocytes into an activated state, known as a cancer-associated adipocyte (CAA) (28-30), resulting in protumorgenic metabolic reprogramming and increased collagen secretion. (31-33) Overall, these findings highlight the plasticity of adipocytes in normal and cancerous mammary glands, yet we have a marginal understanding of the signaling pathways that promote adipocyte plasticity in breast cancer.

To answer these questions, we utilized our immunocompetent *Nf1*-deficient rat model that develops highly penetrant breast cancer. (15) This model contains a germline monoallelic indel in the *Nf1* gene providing a comprehensive approach to study the *Nf1*-deficient stromal contribution to epithelial tumorigenesis. Using several collagen imaging methods, we demonstrated that rats with earlier tumor onset have increased collagen deposition at a younger age. Gene expression analysis revealed that *Nf1* indels increase collagen and ECM gene expression specifically in mammary mature adipocytes before tumor formation. Additionally, these adipocytes also have gene expression consistent with a dedifferentiated preadipocyte/fibroblast-like state. In agreement with our gene expression data, *in vitro* differentiation experiments of rat mammary adipose-derived stem cell (ASC) isolated prior to tumor formation have lower differentiation capacity. Flow cytometry analysis identified heterogeneous adipocyte populations, but no statistically significant changes in ASCs before tumor formation. Therefore, our findings demonstrate that *Nf1* alters adipocyte differentiation and promotes a protumorgenic environment.

## METHODS

### Animals

Female and male immunocompetent *Nf1*-deficient Sprague Dawley rats were bred in-house at the Van Andel Research Institute. The four genotypes include WT, and three *Nf1*-deficient lines: IF/+, PS-20/+, and IF;PS-21/+. Male *Nf1*-deficient rats were bred to wild-type female rats and female pups were used at the indicated developmental time-point. Animals were housed 2 to a cage, had *ad libitum* access to food and water, and were kept under a 12 hr light - 12 hr dark cycle. All animal protocols were approved by the Van Andel Research Institute Animal Care and Use Committee.

### Tissue Histological Processing

Mammary tissues were formalin fixed and paraffin embedded utilizing routine histological procedures. Sections for H&E staining were cut at 5 μm thickness and was performed by Van Andel Institute Pathology and Biorepository Core using a Leica Rotary microtome. Staining was performed on the Tissue-Tek Prisma Plus autostainer, using the Sakura H&E Kit #1. Slides were scanned at 20X magnification on either a Leica Aperio XT or a Leica Aperio AT2 scanning system by the Van Andel Institute Pathology and Biorepository Core.

### H&E Collagen Area Analysis

The H&E collagen area analysis was conducted by Van Andel Institute Optical Imaging Core. Full resolution svs images of H&E-stained mammary tissue sections were imported into QuPath (version 0.4.2) for analysis (34). QuPath’s Estimate Stain Vector tool was used to automatically deconvolve the H&E stain color vectors for each section. A region of interest (ROI) contouring the mammary gland and excluding large blood vessels and lymph nodes was manually generated for each tissue section. A pixel classifier was then created using QuPath’s Pixel Classifier Tool to measure the collagen within the mammary gland ROI. The pixel classifier used an eosin stain-based threshold of 0.05, pixel resolution of 1.0062 um (equivalent to a 2 factor down sample), and Gaussian blur of 1.5; positive pixels were converted into an area measurement within QuPath. The areas of the mammary gland contours and pixel classifier measurements were exported into a csv file for further statistical analysis.

### Picrosirius Red (PSR) Staining

Picrosirius red (PSR) staining was conducted by Van Andel Institute Pathology and Biorepository Core using a standard protocol (35). Briefly, tissue sections 5-μm-thick tissue sections were deparaffinized, treated with phosphomolybdic acid before staining with Sirius red. Slides were washed in hydrochloric acid and 70% EtOH before coverslipping for polarized light imaging.

### Polarized Light Microscopy

Polarized light images were acquired by Van Andel Institute Optical Imaging Core on a Zeiss AxioScan 7, using an EC Plan-Neofluar 20x/0.5 N.A. Pol M27 air objective, and ZEN Blue software (version 3.7). Tissues were illuminated with a Colibri 7 TL LED light source, imaged in brightfield at 5% intensity and 200 μs exposure, and by circularly polarized light at 10% intensity and 3.78 ms exposure, sequentially. Samples were detected by an AxioCam705 color CMOS camera with 5-megapixel resolution. Resulting images were 24 bits and at 0.173x0.173 um scaling per pixel.

### Picrosirius Red Polarized Light Analysis

Full resolution czi images of polarized light collected from picrosirius red-stained mammary tissue sections were imported into QuPath for analysis by Van Andel Institute Optical Imaging Core. A ROI contouring the full tissue section was manually drawn for each slide. A pixel classifier was created using QuPath’s Pixel Classifier Tool to measure the area covered by the signal from polarized light within the ROI. A training image was created from 12 equal sized rectangular regions from 3 different sections using QuPath’s create training image tool. This resulting training image was annotated for examples of signals from the polarized light (n = 53) and background (n = 33) and used for training the pixel classifier using an artificial neural network at 1.38 μm/pixel resolution. All other settings in the pixel classifier were left at their default options. The trained pixel classifier was applied to the ROIs of all images to segment the total area covered by the polarized light signal. The segmented areas were exported from QuPath into a csv file for further statistical analysis.

### Mammary Fat Pad Digestion

Freshly isolated fourth mammary fat pads (MFP) were digested into a single cell suspension as described in Dischinger *et al.* (2018) and Tovar *et al.* (2020). Briefly, finely minced MFP was digested overnight at 37°C in EpiCult B media containing 5% FBS, 1% gentle collagenase/ hyaluronidase, 10 ng/mL recombinant human epidermal growth factor, 10 ng/mL recombinant human basic fibroblast growth factor, and 4 μg/mL heparin. Pelleted cells were incubated with ammonium chloride in HBSS + 2% FBS to lyse red blood cells followed by 0.05% trypsin, and then 5 U/mL dispase + 0.1mg/mL DNase I to further digest cell clumps. Cells were then strained through a 40-μm cell strainer.

### RNA Extraction

For RNA extraction of whole fat pads and tumors, fresh harvested samples were cut up and added to MPbio lysing matrix E tube. Cells were homogenized for 20 seconds in an MPbio homogenizer, then RNA was isolated from the supernatant using the Trizol-chloroform-isopropanol RNA extraction method. For RNA extraction of mature adipocytes, RNA was isolated using the Trizol-chloroform-isopropanol RNA extraction method. For RNA extraction from cultured epithelial cells and fibroblast, 1.2×10^5^ cells were plated into one well of a six-well plate. At 90-100% confluency, RNA was isolated using QIAGEN QIAshredder (QIAGEN #79654) and RNeasy Kit (QIAGEN #74104) manufacturer’s instructions.

### Construction and Sequencing of Directional mRNA-seq Libraries

Libraries were prepared by the Van Andel Institute Genomics Core from 500 ng of total RNA using the KAPA mRNA Hyperprep kit (v4.17) (Kapa Biosystems, Wilmington, MA USA). RNA was sheared to 300-400 bp. Prior to PCR amplification, cDNA fragments were ligated to IDT for Illumina TruSeq UD Indexed adapters (Illumina Inc, San Diego CA, USA). Quality and quantity of the finished libraries were assessed using a combination of Agilent DNA High Sensitivity chip (Agilent Technologies, Inc.), QuantiFluor® dsDNA System (Promega Corp., Madison, WI, USA), and Kapa Illumina Library Quantification qPCR assays (Kapa Biosystems). Individually indexed libraries were pooled and 50 bp, paired-end sequencing was performed on an Illumina NovaSeq6000 sequencer using an S2 sequencing kit (Illumina Inc., San Diego, CA, USA) to an average depth of 45M reads per sample. Base calling was done by Illumina RTA3 and output of NCS was demultiplexed and converted to FastQ format with Illumina Bcl2fastq v1.9.0.

### RNA Sequencing Data Analysis

Raw reads were first trimmed of adapters with Trim_Galore (https://www.bioinformatics.babraham.ac.uk/projects/trim_galore/_) and then mapped to Rnor 6.0 assembly with STAR v2.5.2b using options ‘--twopassMode Basic’ and ‘--quantMode GeneCounts’ to directly output counts for all features from the ensemble annotation 6.0.90 (36). A negative binomial generalized log-linear model was then fit to the filtered count data with edgeR using the weighted trimmed mean of M-values to normalize for library size and composition biases (37). Different genotype or tissue groups were contrasted and p-values were generated using empirical Bayes quasi-likelihood F-tests, and then adjusted using the BH method; adjusted P-values less than 0.05 were considered significant. Heatmaps were generated from library-size normalized counts centered across genes (z-scores) using the R package ComplexHeatmap (38). Using the normalized counts per million, the fold change against matched genotype normal mammary gland expression was calculated for tumor samples and shown as log2 (fold change) capped at -3 and 3.

### Gene Ontology (GO) and Gene Set Enrichment Analysis (GSEA)

The gene ontology (GO) functional enrichment analysis was performed using g:Profiler (version e106_eg53_p16_65fcd97) with g:SCS multiple testing correction method applying significance threshold of *q* < 0.05 (39). Results were plotted using ggplot2 (v3.3.6) (40). For GSEA, protein-coding genes were first pre-ranked by fold change of mutant vs WT and loaded onto GSEA v4.3.2 (41). Analysis was performed on adipocyte differentiation gene signatures obtained from the MSigDB database (42, 43).

### SVF And ASC Isolation

SVF isolation was adapted from Picon-Ruiz *et al.* (2020). After mammary fat pad digestion and red blood cell removal as described above, the SVF fraction was washed with PBS, filtered through a 70-μm cell strainer, resuspended, and plated at 3×10^5^ cells/cm^2^. After 2-3 passages in DMEM supplemented with 10% FBS and 1% penicillin/streptomycin, the cells remaining in culture were taken as the ASC population.

### ASC Differentiation

Adipogenic differentiation was induced using Human MesenCult™ Adipogenic Differentiation Kit (STEMCELL Technologies #05412). According to manufacturer’s protocol, DMEM media was replaced with differentiation media once the ASC culture reached 90-100% confluency. ASC culture was differentiated for 14 days with media changes every 3 days before they were used for downstream applications.

### Oil Red O staining

After 14 days in adipogenic differentiation media, cells were stained with Oil Red O following a standard histological protocol (44). Briefly, cells were fixed with 4% paraformaldehyde for 1 hour at room temperature, then incubated with 60% isopropanol for 5 minutes before staining with Oil Red O (Sigma-Aldrich #O0625) for 30 minutes. After staining, cells were washed with distilled water, stained with Hematoxylin (Vector Laboratories #H-3401) for 1 minute and finally washed with tap water for bluing before visualization/imaging.

### Flow Cytometry Analysis and Sorting of Mammary SVF

For flow cytometry analysis, 1x10^6^ cells in 100 μL were stained for 30 minutes at room temperature using the following antibodies: CD90-BUV395 (BD, #740260, 0.1 mg/μL), ECAD-AF405 (Novus Bio, #NBP2-98433AF405, 1.74 mg/μL), CD45-BV605 (BD, #740371, 0.4 mg/μL), CD29-SB702 (Thermo, #67-0291-82, 0.1 mg/μL), CD34 (Thermo, #PA5-47849, 1 mg/μL) with secondary AF488 (Thermo, #A-11015, 1 mg/μL), CD24-PE (Thermo, #562104, 0.2 mg/μL), CD54-R718 (BD, #751649, 0.1 mg/μL), CD140a-APC (BD, #568234, 0.2 mg/μL), and CD31-PECy7 (Thermo, #25-0310-82, 0.025 mg/μL). All fluorescence minus one (FMO) and single stain controls were stained in HF buffer (HBBS 1X with calcium chloride and magnesium chloride, supplemented with 2% FBS and 1% Pen/Step). Full panel staining was done with HF buffer and 10 μL Brilliant Stain Buffer Plus (BD Horizon #566385) per sample. Finally, cells were washed, stained with 0.005 μg/mL DAPI, filtered, and acquired on the Cytek Biosciences Aurora spectral flow cytometer by the Van Andel Institute Flow Cytometry Core. Data was analyzed using FlowJo v10.8.1. For *in vitro* phenotypic validation, cells were similarly stained and then sorted on the BD FACSymphony S6. Sorted cells were cultured in DMEM with 10% FBS and 1% Pen/Strep.

### Statistical Analysis

#### H&E images eosin quantification

For the day 21 data, we used a mixed-effects beta regression model from the glmmTMB package (45) adjusted for stainer and the scanner types. We also included a random intercept for each unique image. Comparisons between genotypes were performed using emmeans (46). Comparisons are given on the log-odds ratio scale or odds ratio scale and have been adjusted for multiple comparisons using the *dunnettx* method. For the day 33 data, we used a mixed-effects beta regression model from the glmmTMB package (45) that included a random intercept for each unique image. The model was adjusted for the scanner type. There was perfect separation of the stainer and scanner types in the day 33 data, so only one of these variables was used in our models. Comparisons between genotypes were performed using emmeans (46). Comparisons are given on the log-odds ratio scale or odds ratio scale and have been adjusted for multiple comparisons using the *dunnettx* method. All analyses were conducted using R v4.1.0 (https://cran.r-project.org/) with an assumed level of significant of α = 0.05.

#### PSR-POL light quantification

A beta regression model adjusted using Tukey method for multiple comparisons was used to estimate percent of polarized light signals in all samples. To compare percent polarized light in *Nf1*-deficient genotypes to the WT, we used emmeans (46). Results are given on the log odds ratio scale.

#### ASC Differentiation

We used negative binomial mixed effects models with random intercepts for each animal, to estimate and compare the number of cells with lipid droplets against the WT. Tests are performed on the log scale and results are reported on the odds ratio scale.

## RESULTS

### *Nf1*-deficieny Results in Increased Matrix Deposition Before Mammary Tumor Initiation

To investigate the role of *Nf1* in the mammary tumorigenesis, we previously generated three mutant lines representing distinct states of *Nf1* deficiency (Figure 1A). (15) In this model, we targeted the GAP-related domain (GRD) to study the impact of RAS regulation. The IF/+ animals harbor an in-frame indel in exon 20, whereas the PS/+ rats have a premature stop codon in exon 20. The IF;PS-21/+ line was derived from an in-frame indel in exon 20 and a distinct mRNA isoform with a premature stop in exon 21. By creating indels in the GRD of one copy of the *Nf1* gene, each of these *Nf1*-deficient immunocompetent rat lines developed highly penetrant, aggressive mammary adenocarcinomas with varying times on tumor onset. The IF/+ line, with the least impact on neurofibromin function, had the slowest tumor onset, the PS-20/+ and IF;PS- 21/+ lines with a premature stop indel (thus removing all the domains after the GRD) resulted in animals with quicker tumor onset (Figure 1B).

**FIGURE 1:**
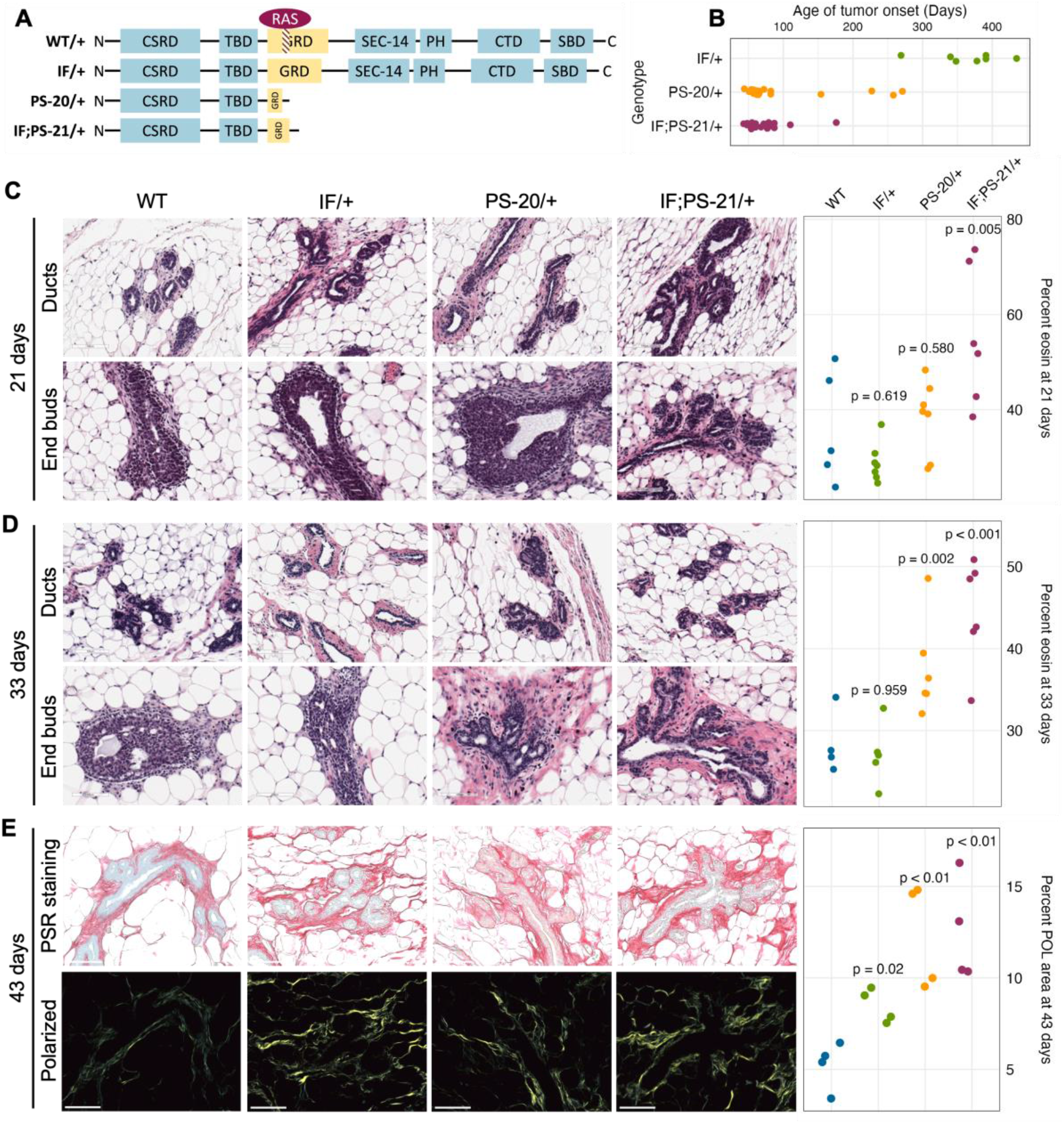
*Nf1*-deficient rat breast cancer lines have increased matrix before tumor formation. (A) Schematic of indels generated in one copy of the *Nf1* gene in immunocompetent Sprague Dawley rats resulting in three *Nf1* mutant lines: *Nf1* in frame (IF/+), premature stop-20 (PS-20/+), as well as in-frame and premature stop-21 (IF;PS-21/+). (B) Tumor onset in *Nf1*- deficient rats. All animals in this study died of mammary tumor progression. Representative H&E images and eosin quantification of rat mammary gland at (A) 21 and (B) 33 days. (C) Representative brightfield and polarized microscopy images of picrosirius red (PSR)-stained rat mammary glands at 43 days, and quantification of polarized light signal. Each data point represents a tissue from an individual rat as a biological replicate.

To understand the mechanism of different tumor onset, it is vital to understand the changes that occur in preneoplastic glands that create an environment of tumor permissiveness. While we have previously shown an increase in mammary epithelial branching and end bud formation during development (47), H&E staining also shows an increased matrix region around the mammary ducts and end buds of *Nf1* deficient rats compared to WT. To quantify this difference, we measured the percent area of eosin around the ducts. At 21 days, the odds of having eosin staining are 14 times higher in *Nf1*-deficient IF;PS-21/+ tissue than WT tissues (p = 0.005, Figure 1C). However, at 33 days, the odds of having eosin staining in *Nf1*-deficient PS- 20/+ and IF;PS-21/+ tissues are 5 times (p = 0.002) and 8 times (p < 0.001) higher respectively compared to WT tissues (Figure 1D). At both 21 and 33 days, the odds of having more eosin staining in *Nf1*-deficient IF/+ tissues than WT tissues are not statistically significant with the odds of having more eosin staining only 2 times higher odds (p = 0.619 at 21 days; p = 0.959 at 33 days).

To verify that this increase in eosin correlates with an increase in collagen deposition that progresses with age towards tumor formation, we stained tissues from 43-day-old rats with picrosirius red (PSR) that allows for collagen visualization under exposure of polarized light. (48) At 43 days, all *Nf1*-deficient mammary glands had increased collagen content compared to the WT glands. Consistent with the eosin quantification, both *Nf1*-deficient PS-20/+ and IF;PS- 21/+ tissues showed approximately 150% increase in polarized light (POL) signals compared to the WT tissues (p < 0.01; Figure 1E). Interestingly, although still lower than the other two *Nf1*-deficient genotypes, the IF/+ tissues at 43 days also had an increase in POL signals as well, about 72% more than the WT tissues (p = 0.02; Figure 1E). This increase in collagen indicates that as the IF/+ rats approach an age closer to tumor formation, their collagen content also increases.

This indicates that collagen and matrix changes correspond to *Nf1* loss of function and rates of tumor onset. In our model, increases in collagen deposition are present earlier in PS-20/+ and IF;PS-21/+ mammary glands which have a larger loss of neurofibromin and earlier tumor onset (6-12 weeks). In contrast, increase in collagen changes are first observed at 43 days, in IF/+ mammary glands that have a minor alteration in neurofibromin and a much later tumor onset (<10 months). These results also show that the timing of increase in collagen mirrors the timing of tumor onset in the *Nf1*-deficient lines.

### *Nf1*-deficient IF/+ Mammary Mature Adipocytes Have Increase Collagen and ECM Genes Expression Before Tumor Formation

To identify which cell population and what type of collagen was predominantly involved in the elevated collagen levels, we performed RNAseq of adipocytes, fibroblasts and epithelial cells isolated from mammary glands before tumor formation. Unsupervised clustering of collagen gene expression revealed that each sample expresses different types of collagens with very little overlap between each other (Supp Fig 1). As expected, fibroblasts have the highest expression of fibrillar collagens, such as collagen I, III, and V, yet there were no significant differences between the *Nf1*-deficient and WT fibroblasts. Epithelial cells and mature adipocytes primarily express distinct subunits of network forming collagens. Surprisingly, we observed that the *Nf1*-deficient IF/+ mature adipocytes have increased gene expression of collagens (Supp Fig 1, Figure 2A). Because adipocytes are more well known for their endocrine and metabolic functions, we were intrigued by these unexpected findings.

**FIGURE 2:**
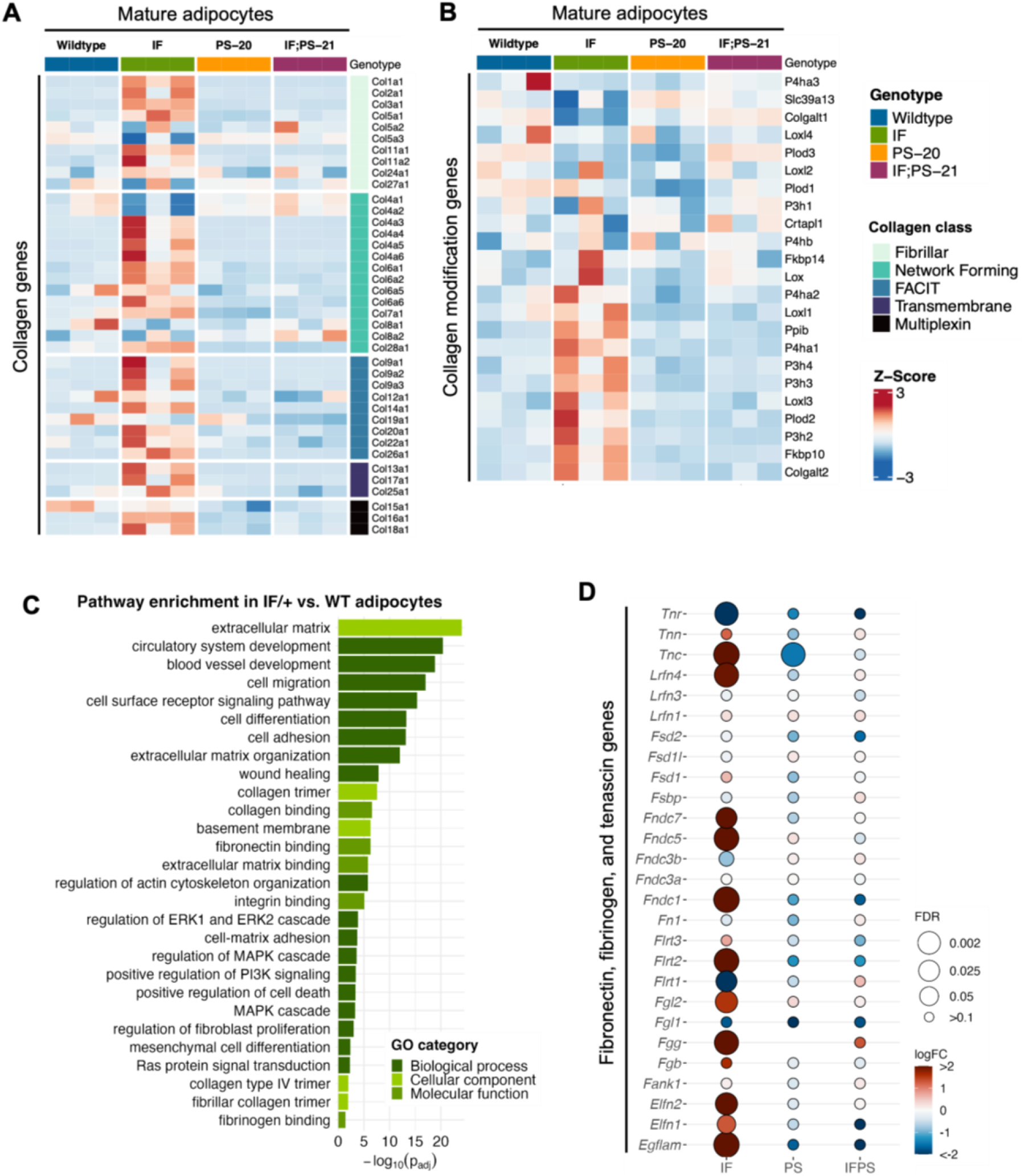
Collagen and ECM gene expression changes in mature adipocytes before tumor formation. Heatmap showing mature adipocyte expression of (A) collagen and (B) collagen modification genes in 33-day old mature adipocytes. (C) Gene ontology (GO) pathway enrichment in IF/+ vs. WT mature adipocytes. (D) Fold change in expression of fibronectin, fibrinogen, and tenascin genes in *Nf1*-deficient mature adipocytes compared to WT.

Recent studies have revealed that adipocytes surrounding a tumor can become activated to form cancer-associated adipocytes (CAAs) and have increased collagen secretion. (27-30, 33) Since the adipocytes we analyzed are from mammary glands before tumor formation, these results indicate that *Nf1* plays a role in normal mammary adipocyte differentiation. Next, we examined whether the IF/+ mature adipocytes also exhibit other reported gene expression features of CAAs. Along with increased collagen secretion, these studies also reported that CAAs can remodel collagen architecture and alignment through increased expression of collagen modification genes. (33) An example of a collagen modification gene is prolyl 4-hydroxylase subunit alpha 1 (*P4ha1*), which is the most important catalytic subunit of the P4H enzyme and necessary for collagen polypeptide chains folding into stable triple-helical molecules (49). From our RNAseq data, *P4ha1* and other collagen modification genes have increased expression in IF/+ mature adipocytes compared to the WT and two other *Nf1*-deficient mature adipocytes (Figure 2B). Additionally, gene ontology (GO) analysis of *Nf1*-deficient IF/+ mature adipocytes gene expression compared to WT showed enrichment in pathways involved in ECM processes and RAS-MAPK signaling (Figure 2C). Pathways involved in MAPK signaling were expected due to neurofibromin’s role as the main negative regulator of RAS. Enrichment of pathways in ECM processes substantiates a role for *Nf1*-deficient IF/+ adipocytes in collagen deposition and a CAA-like phenotypes (Figure 2A-B). Our analysis indicated other ECM components besides collagen may contribute to tumorigenesis in *Nf1*-deficient IF/+ glands. Several genes typically known for their role in tissue fibrosis (i.e., tenascin, fibrinogen, and fibronectin genes) were increased in *Nf1*-deficient IF/+ adipocytes (Figure 3D). Overall, these results demonstrated that *Nf1*-deficient IF/+ adipocytes have increased collagen and fibrosis-related ECM gene expression preceding tumor formation.

**FIGURE 3:**
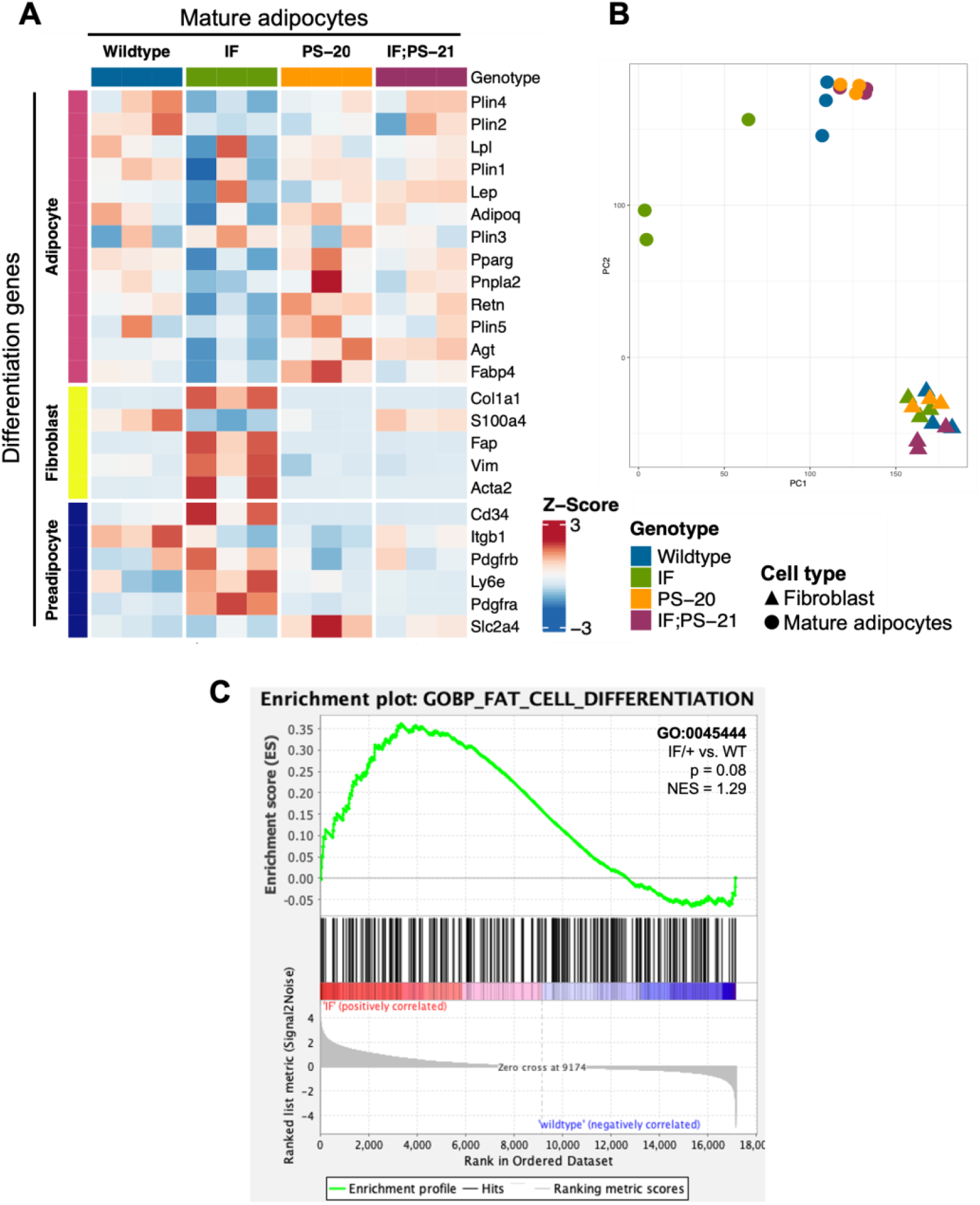
*Nf1*-deficient IF/+ mammary mature adipocytes before tumor formation mimic a preadipocyte and fibroblast-like cell state. (A) Heatmap showing mesenchymal lineage differentiation genes expression in 33-day old mature adipocytes. (B) PCA plot of fibroblasts and adipocytes. (C) Enrichment plot of fat cell differentiation gene set (GO:0045444, p = 0.08, NES = 1.29) in IF/+ vs. WT mature adipocytes.

### *Nf1*-Deficient IF/+ Mammary Adipocytes Mimic a Preadipocyte and Fibroblast-Like Cell State

Another reported phenotype of CAAs is their ability to dedifferentiate to a preadipocyte- and fibroblast-like cell through a process called adipocyte mesenchymal transition (AMT). (27-30) To determine if AMT was present in the *Nf1*-deficient IF/+ mature adipocytes, we evaluated adipocyte differentiation expression signatures. Principal component analysis (PCA) of adipocyte and fibroblast samples show that the *Nf1*-deficient IF/+ mature adipocytes cluster further away from the other *Nf1*-deficient genotypes, indicating that their gene expression profile has deviated from a typical mature adipocyte (Figure 3B). Looking at the different genes expressed at adipocyte, preadipocyte, and fibroblast stages, our RNAseq data showed the *Nf1*-deficient IF/+ mature adipocytes had lower expression of mature adipocyte markers, but higher expression of fibroblast and preadipocyte markers. WT and *Nf1*-deficient PS-20/+ and IF;PS-21/+ mature adipocytes do not have altered mesenchymal lineage gene expression. For example, adiponectin (*Adipoq*) and perilipin-1 (*Plin1*), genes which are primarily expressed in adipose tissues and mature adipocytes but are both downregulated in the IF/+ mature adipocytes. In contrast, fibroblast activation protein alpha (*Fap*) and *Cd34* are genes primarily expressed in activated fibroblast and preadipocytes, respectively, but have increased expression in the IF/+ mature adipocytes compared to the other genotypes (Figure 3A). Gene set enrichment analysis (GSEA) of the mature adipocytes revealed an enrichment in adipocyte differentiation gene signatures (obtained from the MSigDB database). To highlight one example, the *Nf1*-deficient mature adipocytes showed an enrichment in a fat cell differentiation gene set (GO0045444) which supports the difference in mesenchymal lineage markers we observed (NES = 1.29, p = 0.08; Figure 3C). Overall, these results show that the IF/+ indel in one *Nf1* allele causes the gene expression profile of mature adipocytes before tumor formation to mimic a preadipocyte- and fibroblast-like cell state.

### *Nf1* Deficiency Restricts Mammary Adipose-Derived Stem Cell Differentiation to Immature Adipocytes

To verify the altered differentiation gene expression observed in our RNAseq data (Figure 3), we isolated the adipose-derived stem cell (ASC) population from our rat mammary glands and differentiated them *in vitro* as illustrated in Figure 4A. After an overnight collagenase digest, we cultured the stromal vascular fraction (SVF) for 2-3 passages to enrich for the ASC population before culturing them in commercially available differentiation media for 2 weeks. After 2 weeks, fully differentiated cells containing lipid droplets were counted as a measure of differentiation ability. After 14 days in differentiation media, we observed that *Nf1*-deficient IF/+ ASCs have 60% fewer cells with lipid droplets compared to WT, thus indicating a decreased differentiation ability (WT vs. IF/+ odds ratio = 0.4, p = 0.08; Figure 4C).

**FIGURE 4:**
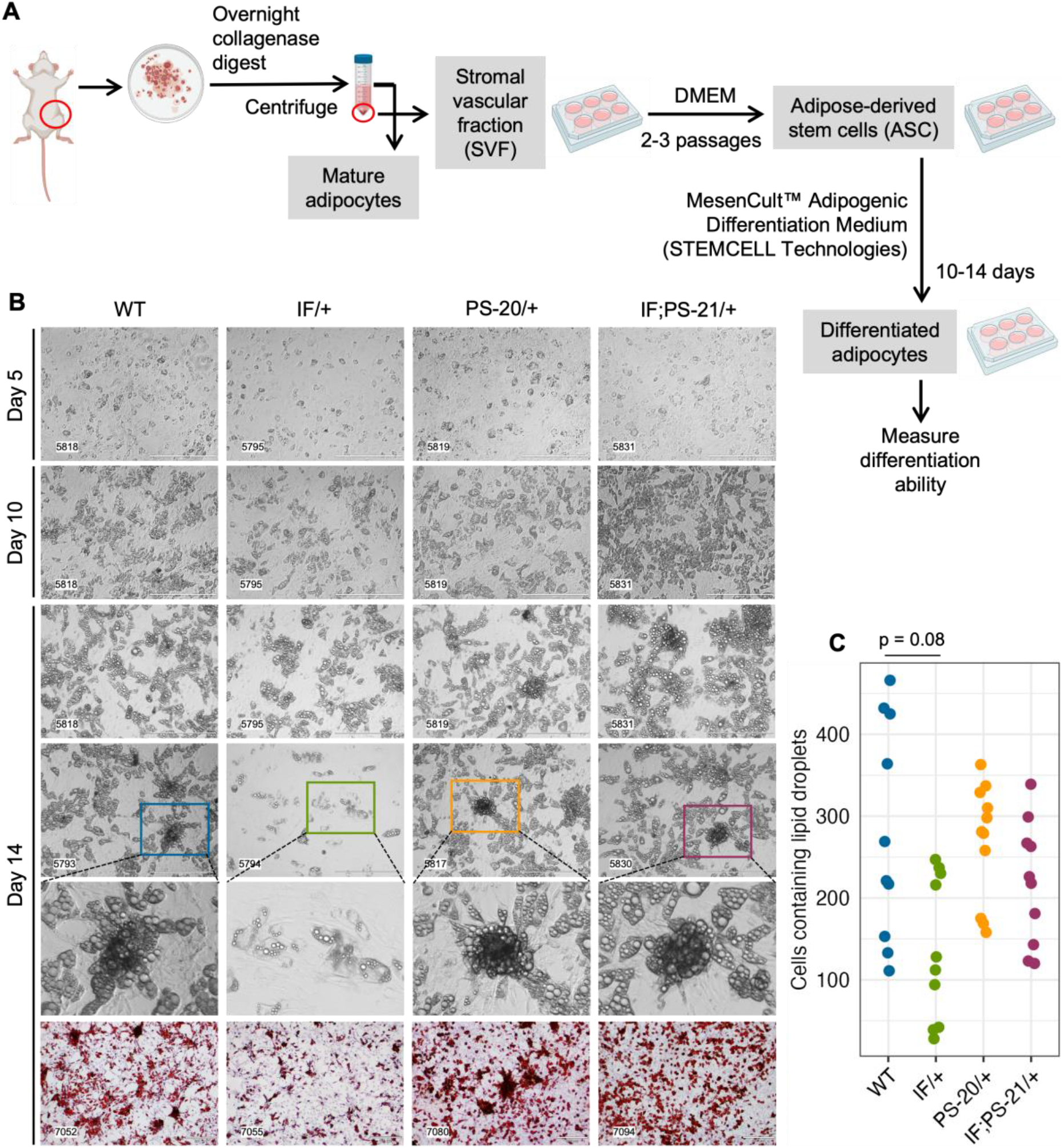
*Nf1* deficiency restricts mammary adipose-derived stem cells (ASCs) differentiation to immature adipocytes. (A) Workflow of isolating and differentiating stromal vascular fraction (SVF) into ASCs and differentiating them further into mature adipocytes. (B) Brightfield images of ASC differentiation at day 5, 10, and 14, as well as Oil Red O staining at day 14. Brightfield images were acquired at 10X magnification, scale bar is 400 µm. Oil Red O images were acquired at 4X magnification, scale bar is 300 µm. (C) Quantification of cells containing lipid droplets at day 14 (WT vs IF/+ odds ratio = 2.39, p = 0.08; n =3 rats per genotype).

### *Nf1* Deficiency Does Not Decrease Adipose-Derived Stem Cell Populations in the Mammary Stromal Vascular Fraction

Mature adipocytes and preadipocytes differentiate through a multistep process originating from mesenchymal stem cells (MSCs), and in adipose tissues specifically, these tissue resident MSCs are known as ASCs. (50-52) To investigate whether the reduced differentiation ability of the *Nf1*-deficient IF/+ ASC that we observed (Figure 4) was due to a decreased ASC population, we used flow cytometry to quantify the amount of ASC present in the SVF of our rat mammary glands (Figure 5A). After red blood cell removal, we derived a lineage-negative population by eliminating CD45+ hematopoietic cells, CD31+ endothelial cells, and E-Cad+ epithelial cells.

**FIGURE 5:**
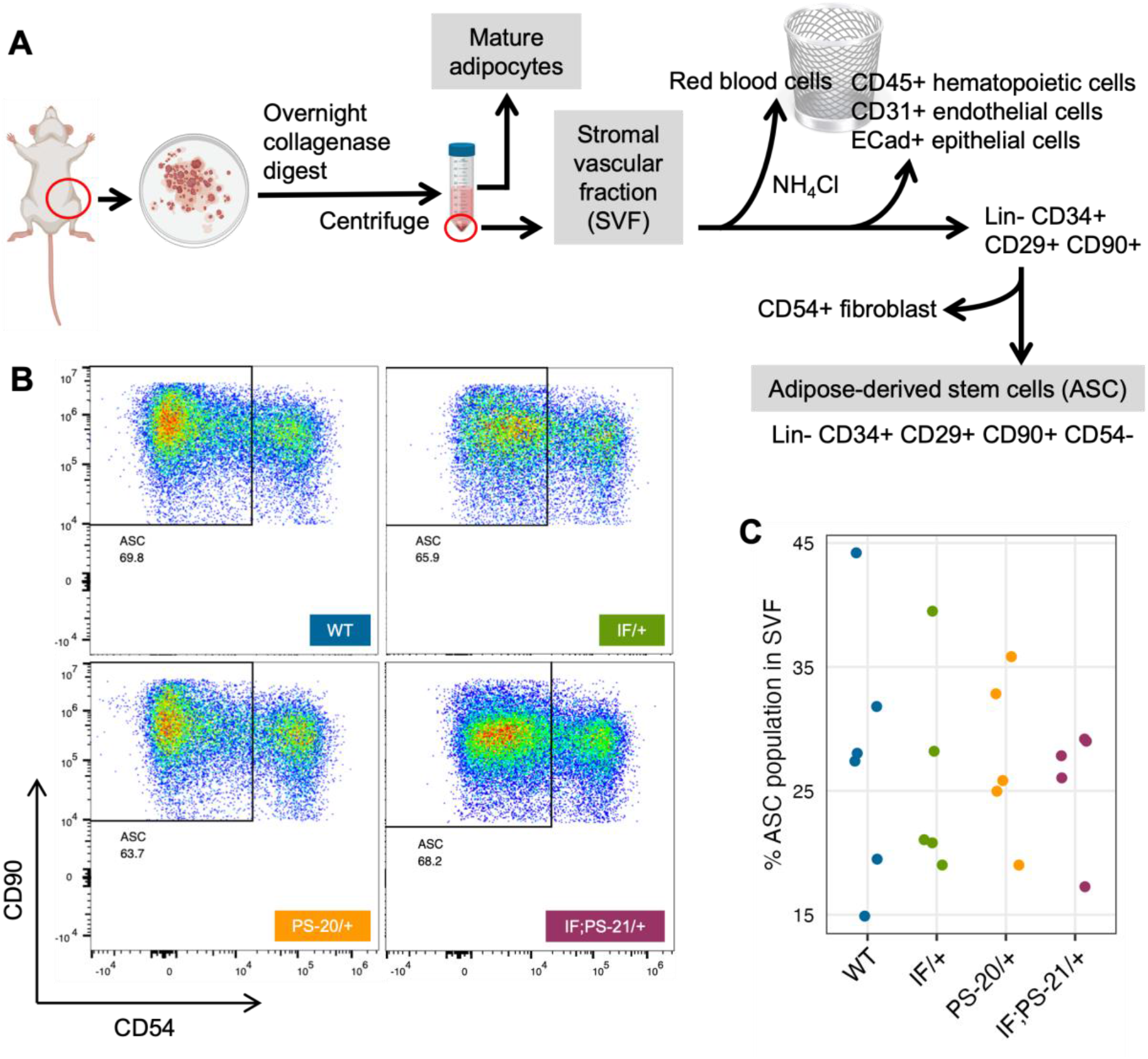
*Nf1* deficiency does not decrease adipose-derived stem cell (ASC) population in the mammary stromal vascular fraction (SVF). (A) Workflow of isolating ASC population from 33-day old rat mammary gland SVF using FACS. (B) Flow cytometry analysis of Lin-CD34+ CD29+ CD90+ CD54-rat mammary gland ASC. (C) Percent ASC population present in pooled SVF population.

Using three stem cell markers, CD34+ CD29+ CD90+, we further stratified the ASC population, before excluding CD54+ fibroblasts. (50, 51, 53-55) The abundance of the resulting ASC population (Lin-CD34+ CD29+ CD90+ CD54-) was measured compared to the whole SVF population. Even though substantial heterogeneity of ASC was observed within the genotypes, we did not detect a decrease of the Lin-CD34+ CD29+ CD90+ CD54-ASC population in *Nf1*-deficient IF/+ rats compared to WT (Figure 5C). Together, our differentiation assays and flow cytometry analysis indicate that *Nf1*-deficient IF/+ adipocytes have impaired differentiation that is not due to a decrease in the adipocyte progenitor population. In addition to demonstrating that *Nf1* does not impact the early stages of differentiation, these data suggest a potential of *Nf1* regulating the later stages of adipocyte differentiation instead, specifically through PPARγ signaling.

## DISCUSSION

Recent breast cancer studies have highlighted the important role that *NF1* dysregulation has in both *NF1*-related and sporadic breast cancers with the focus being on epithelial cells. (47, 56) With the goal of uncovering non-epithelial contributions to NF1-related breast cancer tumorigenesis, we measured alterations in the *Nf1*-deficient mammary stromal region before tumor formation. Here we show for the first time that *Nf1*-deficiency impacts mammary stroma and the extracellular matrix, especially collagen formation, before mammary tumor formation. In the field of NF1 research, studies evaluating collagen in NF1-mediated tumorigenesis have only been conducted in the context of neurofibromas and nerve sheath tumors. (57-59) Despite the known impact of collagen in breast cancer as well as other NF1-related tumors, the contribution of collagen in NF1-associated breast cancer has not been characterized. The findings in the study demonstrate that *Nf1*-deficient rats with the earliest tumor onset have a significant increase in collagen expression compared to the *Nf1*-deficient rats with later tumor onset. Notably, these collagen and matrix changes were observed weeks before tumor formation typically occurs in the these *Nf1*-deficient glands. These results indicate that increased collagen deposition by *Nf1*-deficient stromal cells is associated with earlier tumor development.

Next, we evaluated which stromal populations were driving altered matrix deposition in the pretumorigenic Nf1-deficient mammary glands. Our initial assumption was that fibroblasts were the predominant cell responsible for increasing matrix deposition, but RNAseq data led us to focus on the mammary adipocytes. Interestingly, the RNAseq analysis revealed that *Nf1*-deficient IF/+ adipocytes before tumor formation express more collagen and collagen modification genes, which are commonly associated as a feature of fibroblasts. Pathway enrichment analysis identified pathways involved in MAPK signaling, ECM deposition, and adipocyte differentiation. Additionally, *Nf1*-deficient adipocytes have a fibroblast- and preadipocyte-like expression profile and decreased expression of mature adipocyte markers. To determine if *Nf1* loss was impacting adipocyte differentiation, we isolated ASCs from the SVF population of rat mammary gland before tumor formation and observed decreased differentiation of *Nf1*-deficient IF/+ ASCs; however, flow cytometry analysis demonstrated that the abrogated differentiation was not due to a decrease in the Lin-CD34+ CD29+ CD90+ CD54-ASC population. These results indicate that this *Nf1* loss of function promotes a preadipocyte-like phenotype, but this cell state change does not originate in the ASC population.

In addition to identifying a role for *Nf1* in promoting adipocyte plasticity, these findings also uncover important questions regarding *Nf1* loss in breast cancer initiation. The most crucial question is why AMT only observed in mature adipocytes from *Nf1*-deficient IF/+ that have the slowest tumor onset (compared to PS-20/+ and IF;PS-21/+ *Nf1*-deficient lines). There are several potential explanations for this distinct stromal phenotype. First, we may be missing the short window of the tumor microenvironmental alterations that occur in the rapid tumor onset that occur in *Nf1*-deficient PS-20/+ and IF;PS-21/+ glands. Alternatively, these distinct stromal differences may be due to distinct signaling changes present in the IF/+ indel (only impacting the GRD domain) compared to the premature stop indels in the rapid tumor lines. This difference indicates that *Nf1* has cell-specific functions and potentially different corresponding signaling mechanisms.

While our work and other studies have shown that the epithelial transformation is in part due to the NF1 corepressor function of ERα (56), it is unclear whether NF1-ER signaling impacts adipocyte differentiation to promote a protumorigenic environment. Our *Nf1*-deficient models were created by targeting the GRD domain to alter RAS signaling. (15) There are several studies elucidating the role of RAS in regulating PPARγ activity that can be applicable in the context of *NF1* deficiency (Figure 6A). PPARγ is the master regulator of adipocyte differentiation (60) and is required for mature adipocytes to fully differentiate and maintain their differentiated state. (61-63) The RAS effector ERK has been shown to phosphorylate PPARγ at Ser112 and consequently suppress PPARγ transcriptional activity. (64, 65) Concurrently, inhibition of KRAS has also been shown to promote *in vitro* adipogenic differentiation in the 3T3-L1 and C2C12 preadipocyte cell lines. (66) Alternatively, RAS can indirectly suppress PPARγ activity through TAZ, the main downstream effector protein of the Hippo pathway. In its active state, TAZ is a transcriptional regulator that binds to PPARγ and prevents DNA binding; but in its inactive state, TAZ is phosphorylated and undergoes proteasomal degradation. (67) RAS can prevent phosphorylation and degradation of TAZ, thus suppressing adipocyte differentiation. (68) Interestingly, a recent study also demonstrated that TAZ binding to PPARγ is increased with ERK-mediated phosphorylation of PPARγ at Ser112. Therefore, loss of NF1 function may impact adipocyte differentiation through RAS, PPARγ, and Hippo signaling alterations.(69) Importnatly, the lack of change in ASC abundance, implies that NF1 may modulate later stages of adipocyte differentiation through PPARγ. Our future studies into these signaling mechanisms will provide vital insights into adipocyte plasticity not only in breast cancer, but also in ovarian cancer which is another cancer that occurs in an adipose-rich organ and is also highly prevalent among NF1 patients. (13)

**FIGURE 6:**
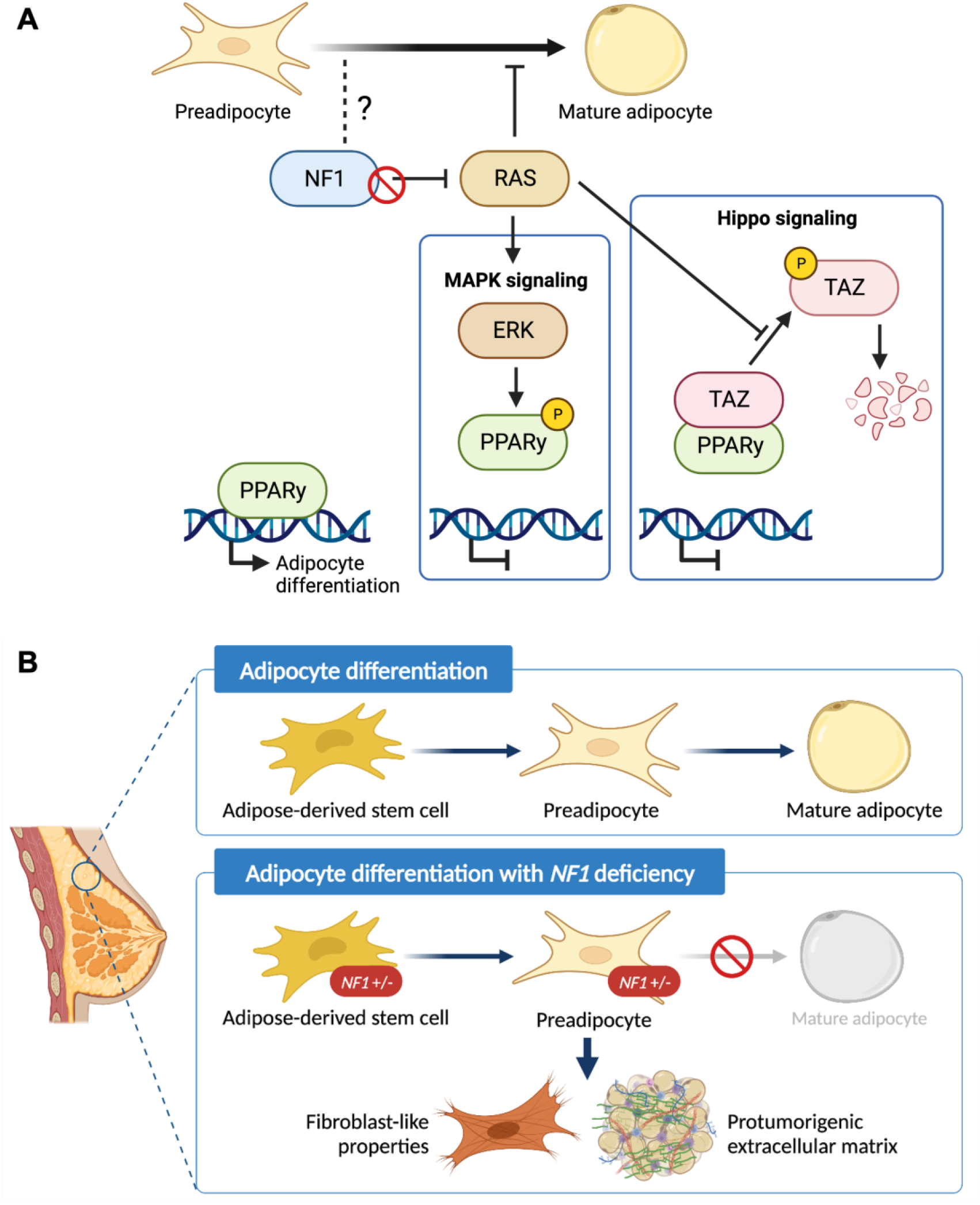
Schematic of the role of *NF1* in mammary adipocyte differentiation. (A) Potential mechanism of how loss of *NF1* suppresses PPARγ signaling during the final stages of adipocyte differentiation through the MAPK and Hippo signaling pathways. (B) Adipose-derived stem cells (ASCs) differentiate into preadipocytes before finally differentiating into mature adipocytes. In the context of *NF1* deficiency, we show that mammary ASCs are restricted to the preadipocyte stage with fibroblast-like properties and can potentially secrete protumorigenic extracellular matrix before tumor formation.

In summary, findings presented in study have uncovered the previously unknown role of *Nf1* in regulating mammary collagen deposition and adipocyte plasticity (Figure 6B). These newly characterized roles of *Nf1* bring us closer to answering the clinically relevant question of why NF1 patients have an increased risk of developing breast cancer with poorer prognosis compared to the general population. Understanding the impact of adipocyte plasticity in breast cancer will allow us to identify potential vulnerabilities that can be leveraged for breast cancer early detection and treatment. (70, 71) Additionally, it also creates exciting new research questions regarding the signaling mechanism of *NF1* in distinct cell types as well as other aspects of adipose-related symptom manifestation in individuals with NF1.

## LIST OF ABBREVIATIONS

NF1: neurofibromatosis type 1
RAS: rat sarcoma virus
GRD: GAP-related domain
MAPK: mitogen-activated protein kinase
ECM: extracellular matrix
CAF: cancer-associated fibroblast
AMT: adipocyte-mesenchymal transition
CAA: cancer-associated adipocyte
ASC: adipose-derived stem cells
SVF: stromal vascular fraction
PSR: picrosirius red staining

## DECLARATIONS

### Ethics approval and consent to participate

All animal protocols were approved by the Van Andel Research Institute Animal Care and Use Committee.

### Consent for publication

Not applicable.

### Availability of data and materials

The raw data of bulk RNA-seq generated by this study have been deposited in NCBI’s Gene Expression Omnibus and are accessible through GEO Series accession number GEO: GSE231603. This paper does not report original code. Any additional information required to reanalyze the data reported in this work paper is available from the corresponding authors upon request.

### Authors’ contributions

MA conceptualized and conducted all the experiments. PSD developed the rat model. EAT and PSD isolated and extracted RNA from different mammary populations for RNAseq. CJE maintain the rat colonies and performed the animal protocols. ITB conducted bioinformatics data analysis of the RNAseq data. EW and ZBM conducted the power calculations and statistical analysis. LT and KF performed the histological processing and scanning. KLG, LC, and CRE contributed at all stages of image acquisition and analysis. MN and RTCS conducted flow cytometry method optimization and data analysis. GM provided expertise on image acquisition. Original manuscript was written by MA and reviewed by CRG and MRS. Funding acquisition, resources, and supervision was provided by MRS and CRG. All authors read and approved the final manuscript.

## Acknowledgements

This work was possible thanks to the excellent service from the Van Andel Institute cores (Vivarium and Transgenics, Pathology and Biorepository, Optical Imaging, Genomics, Flow Cytometry, Bioinformatics and Biostatistics).

## SUPPLEMENTAL FIGURES

**Supplemental Figure 1:**
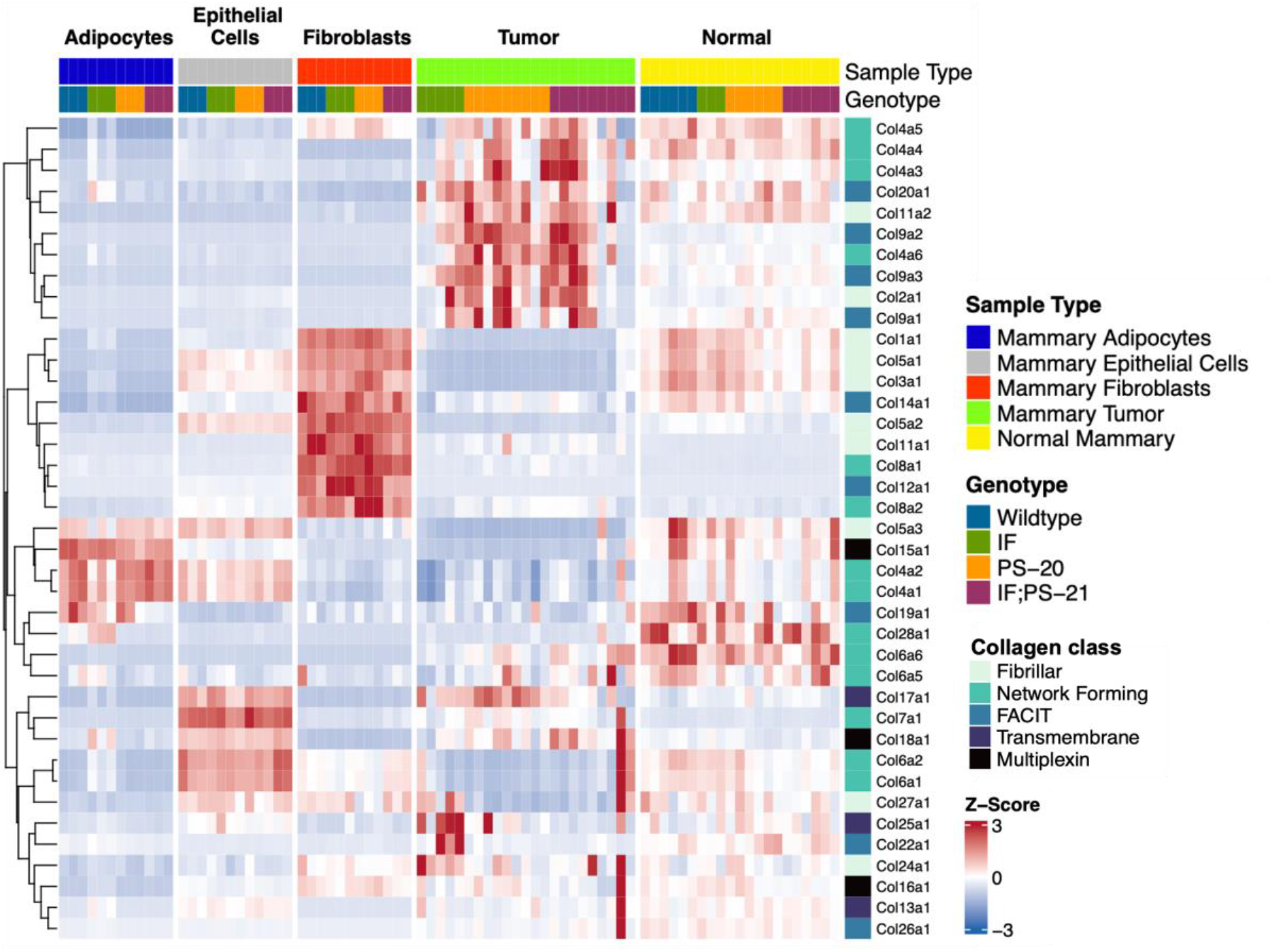
Distinct collagen expression in different cell populations of the mammary gland. Z-scores of collagen gene expression in tumors as well as adipocytes, epithelial cells, fibroblast, whole normal mammary gland before tumor formation at 33 days old.

## REFERENCES

1. Rasmussen SA, Friedman JM. NF1 gene and neurofibromatosis 1. Am J Epidemiol. 2000;151(1):33–40.

2. Griffiths S, Thompson P, Frayling I, Upadhyaya M. Molecular diagnosis of neurofibromatosis type 1: 2 years experience. Fam Cancer. 2007;6(1):21–34.

3. Mo J, Moye SL, McKay RM, Le LQ. Neurofibromin and suppression of tumorigenesis: beyond the GAP. Oncogene. 2022;41(9):1235–51.

4. Yap YS, McPherson JR, Ong CK, Rozen SG, Teh BT, Lee AS, et al. The NF1 gene revisited - from bench to bedside. Oncotarget. 2014;5(15):5873–92.

5. Xu GF, O’Connell P, Viskochil D, Cawthon R, Robertson M, Culver M, et al. The neurofibromatosis type 1 gene encodes a protein related to GAP. Cell. 1990;62(3):599–608.

6. Ratner N, Miller SJ. A RASopathy gene commonly mutated in cancer: the neurofibromatosis type 1 tumour suppressor. Nat Rev Cancer. 2015;15(5):290–301.

7. Sharif S, Moran A, Huson SM, Iddenden R, Shenton A, Howard E, et al. Women with neurofibromatosis 1 are at a moderately increased risk of developing breast cancer and should be considered for early screening. J Med Genet. 2007;44(8):481–4.

8. Madanikia SA, Bergner A, Ye X, Blakeley JO. Increased risk of breast cancer in women with NF1. Am J Med Genet A. 2012;158A(12):3056-60.

9. Wang X, Levin AM, Smolinski SE, Vigneau FD, Levin NK, Tainsky MA. Breast cancer and other neoplasms in women with neurofibromatosis type 1: a retrospective review of cases in the Detroit metropolitan area. Am J Med Genet A. 2012;158A(12):3061-4.

10. Seminog OO, Goldacre MJ. Age-specific risk of breast cancer in women with neurofibromatosis type 1. Br J Cancer. 2015;112(9):1546–8.

11. Uusitalo E, Kallionpaa RA, Kurki S, Rantanen M, Pitkaniemi J, Kronqvist P, et al. Breast cancer in neurofibromatosis type 1: overrepresentation of unfavourable prognostic factors. Br J Cancer. 2017;116(2):211–7.

12. Uusitalo E, Rantanen M, Kallionpaa RA, Poyhonen M, Leppavirta J, Yla-Outinen H, et al. Distinctive Cancer Associations in Patients With Neurofibromatosis Type 1. J Clin Oncol. 2016;34(17):1978–86.

13. Landry JP, Schertz KL, Chiang YJ, Bhalla AD, Yi M, Keung EZ, et al. Comparison of Cancer Prevalence in Patients With Neurofibromatosis Type 1 at an Academic Cancer Center vs in the General Population From 1985 to 2020. JAMA Netw Open. 2021;4(3):e210945.

14. Daly MB, Pilarski R, Berry M, Buys SS, Farmer M, Friedman S, et al. NCCN Guidelines Insights: Genetic/Familial High-Risk Assessment: Breast and Ovarian, Version 2.2017. J Natl Compr Canc Netw. 2017;15(1):9-20.

15. Dischinger PS, Tovar EA, Essenburg CJ, Madaj ZB, Gardner EE, Callaghan ME, et al. NF1 deficiency correlates with estrogen receptor signaling and diminished survival in breast cancer. NPJ Breast Cancer. 2018;4:29.

16. Easton DF, Pharoah PD, Antoniou AC, Tischkowitz M, Tavtigian SV, Nathanson KL, et al. Gene-panel sequencing and the prediction of breast-cancer risk. N Engl J Med. 2015;372(23):2243–57.

17. Finak G, Bertos N, Pepin F, Sadekova S, Souleimanova M, Zhao H, et al. Stromal gene expression predicts clinical outcome in breast cancer. Nat Med. 2008;14(5):518–27.

18. Oh EY, Christensen SM, Ghanta S, Jeong JC, Bucur O, Glass B, et al. Extensive rewiring of epithelial-stromal co-expression networks in breast cancer. Genome Biol. 2015;16(1):128.

19. Bainer R, Frankenberger C, Rabe D, An G, Gilad Y, Rosner MR. Gene expression in local stroma reflects breast tumor states and predicts patient outcome. Sci Rep. 2016;6:39240.

20. de Visser KE, Joyce JA. The evolving tumor microenvironment: From cancer initiation to metastatic outgrowth. Cancer Cell. 2023;41(3):374–403.

21. Curran CS, Ponik SM. The Mammary Tumor Microenvironment. 12962020. p. 163–81.

22. Nee K, Ma D, Nguyen QH, Pein M, Pervolarakis N, Insua-Rodriguez J, et al. Preneoplastic stromal cells promote BRCA1-mediated breast tumorigenesis. Nat Genet. 2023;55(4):595–606.

23. Hennighausen L, Robinson GW. Signaling pathways in mammary gland development. Dev Cell. 2001;1(4):467–75.

24. Zwick RK, Rudolph MC, Shook BA, Holtrup B, Roth E, Lei V, et al. Adipocyte hypertrophy and lipid dynamics underlie mammary gland remodeling after lactation. Nat Commun. 2018;9(1):3592.

25. Corsa CAS, MacDougald OA. Cyclical Dedifferentiation and Redifferentiation of Mammary Adipocytes. Cell Metab. 2018;28(2):187–9.

26. Wang QA, Song A, Chen W, Schwalie PC, Zhang F, Vishvanath L, et al. Reversible De-differentiation of Mature White Adipocytes into Preadipocyte-like Precursors during Lactation. Cell Metab. 2018;28(2):282–8 e3.

27. Zhu Q, Zhu Y, Hepler C, Zhang Q, Park J, Gliniak C, et al. Adipocyte mesenchymal transition contributes to mammary tumor progression. Cell Rep. 2022;40(11):111362.

28. Dirat B, Bochet L, Dabek M, Daviaud D, Dauvillier S, Majed B, et al. Cancer-associated adipocytes exhibit an activated phenotype and contribute to breast cancer invasion. Cancer Res. 2011;71(7):2455–65.

29. Zhang Y, Daquinag AC, Amaya-Manzanares F, Sirin O, Tseng C, Kolonin MG. Stromal progenitor cells from endogenous adipose tissue contribute to pericytes and adipocytes that populate the tumor microenvironment. Cancer Res. 2012;72(20):5198–208.

30. Bochet L, Lehuede C, Dauvillier S, Wang YY, Dirat B, Laurent V, et al. Adipocyte-derived fibroblasts promote tumor progression and contribute to the desmoplastic reaction in breast cancer. Cancer Res. 2013;73(18):5657–68.

31. Bi P, Yue F, Karki A, Castro B, Wirbisky SE, Wang C, et al. Notch activation drives adipocyte dedifferentiation and tumorigenic transformation in mice. J Exp Med. 2016;213(10):2019–37.

32. Wang YY, Valet P, Muller C, Wang YY, Attané C, Milhas D, et al. Mammary adipocytes stimulate breast cancer invasion through metabolic remodeling of tumor cells. JCI insight. 2017;2(4):1–21.

33. Wei X, Li S, He J, Du H, Liu Y, Yu W, et al. Tumor-secreted PAI-1 promotes breast cancer metastasis via the induction of adipocyte-derived collagen remodeling. Cell Commun Signal. 2019;17(1):58.

34. Bankhead P, Loughrey MB, Fernandez JA, Dombrowski Y, McArt DG, Dunne PD, et al. QuPath: Open source software for digital pathology image analysis. Sci Rep. 2017;7(1):16878.

35. Dolber PC, Spach MS. Picrosirius red staining of cardiac muscle following phosphomolybdic acid treatment. Stain Technol. 1987;62(1):23–6.

36. Dobin A, Davis CA, Schlesinger F, Drenkow J, Zaleski C, Jha S, et al. STAR: ultrafast universal RNA-seq aligner. Bioinformatics. 2013;29(1):15–21.

37. Robinson MD, McCarthy DJ, Smyth GK. edgeR: a Bioconductor package for differential expression analysis of digital gene expression data. Bioinformatics. 2010;26(1):139–40.

38. Gu Z, Eils R, Schlesner M. Complex heatmaps reveal patterns and correlations in multidimensional genomic data. Bioinformatics. 2016;32(18):2847–9.

39. Raudvere U, Kolberg L, Kuzmin I, Arak T, Adler P, Peterson H, et al. g:Profiler: a web server for functional enrichment analysis and conversions of gene lists (2019 update). Nucleic Acids Res. 2019;47(W1):W191–W8.

40. Wickham H. ggplot2: Elegant Graphics for Data Analysis: Springer-Verlag New York; 2016.

41. Subramanian A, Tamayo P, Mootha VK, Mukherjee S, Ebert BL, Gillette MA, et al. Gene set enrichment analysis: a knowledge-based approach for interpreting genome-wide expression profiles. Proc Natl Acad Sci U S A. 2005;102(43):15545–50.

42. Liberzon A, Birger C, Thorvaldsdottir H, Ghandi M, Mesirov JP, Tamayo P. The Molecular Signatures Database (MSigDB) hallmark gene set collection. Cell Syst. 2015;1(6):417–25.

43. Liberzon A, Subramanian A, Pinchback R, Thorvaldsdottir H, Tamayo P, Mesirov JP. Molecular signatures database (MSigDB) 3.0. Bioinformatics. 2011;27(12):1739–40.

44. Suvarna KL, Christopher; Bancroft, John D. Bancroft’s Theory and Practice of Histological Techniques. Eighth ed: Elsevier; 2019. 557 p.

45. Brooks ME, Kristensen K, Benthem KJv, Magnusson A, Berg CW, Nielsen A, et al. glmmTMB Balances Speed and Flexibility Among Packages for Zero-inflated Generalized Linear Mixed Modeling. The R Journal. 2017;9(2).

46. Lenth RV. emmeans: Estimated Marginal Means, aka Least-Squares Means. R package version 1.8.0. 2022 [Available from: https://CRAN.R-project.org/package=emmeans.

47. Tovar EA, Arumugam M, Essenburg CJ, Dischinger PS, Grit JL, Callaghan ME, et al. Nf1 deficiency accelerates mammary development and promotes luminal-basal plasticity. bioRxiv. 2022:2022.12.22.520633-2022.12.22.

48. Coelho PGB, Souza MV, Conceicao LG, Viloria MIV, Bedoya SAO. Evaluation of dermal collagen stained with picrosirius red and examined under polarized light microscopy. An Bras Dermatol. 2018;93(3):415–8.

49. Myllyharju J, Kivirikko KI. Collagens, modifying enzymes and their mutations in humans, flies and worms. Trends Genet. 2004;20(1):33–43.

50. Berry R, Rodeheffer MS. Characterization of the adipocyte cellular lineage in vivo. Nat Cell Biol. 2013;15(3):302–8.

51. Berry R, Rodeheffer MS, Rosen CJ, Horowitz MC. Adipose Tissue Residing Progenitors (Adipocyte Lineage Progenitors and Adipose Derived Stem Cells (ADSC). Curr Mol Biol Rep. 2015;1(3):101–9.

52. Brown AC. Insights into the adipose stem cell niche in health and disease. In: Kokai L, Marra K, Rubin JP, editors. Scientific Principles of Adipose Stem Cells: Academic Press; 2022. p. 57-80.

53. Rodeheffer MS, Birsoy K, Friedman JM. Identification of white adipocyte progenitor cells in vivo. Cell. 2008;135(2):240–9.

54. Jeffery E, Church CD, Holtrup B, Colman L, Rodeheffer MS. Rapid depot-specific activation of adipocyte precursor cells at the onset of obesity. Nat Cell Biol. 2015;17(4):376–85.

55. Jeffery E, Wing A, Holtrup B, Sebo Z, Kaplan JL, Saavedra-Pena R, et al. The Adipose Tissue Microenvironment Regulates Depot-Specific Adipogenesis in Obesity. Cell Metab. 2016;24(1):142–50.

56. Zheng ZY, Anurag M, Lei JT, Cao J, Singh P, Peng J, et al. Neurofibromin Is an Estrogen Receptor-alpha Transcriptional Co-repressor in Breast Cancer. Cancer Cell. 2020;37(3):387–402 e7.

57. Peltonen J, Penttinen R, Larjava H, Aho HJ. Collagens in neurofibromas and neurofibroma cell cultures. Ann N Y Acad Sci. 1986;486:260–70.

58. Brosseau JP, Sathe AA, Wang Y, Nguyen T, Glass DA, 2nd, Xing C, et al. Human cutaneous neurofibroma matrisome revealed by single-cell RNA sequencing. Acta Neuropathol Commun. 2021;9(1):11.

59. Atit RP, Crowe MJ, Greenhalgh DG, Wenstrup RJ, Ratner N. The Nf1 tumor suppressor regulates mouse skin wound healing, fibroblast proliferation, and collagen deposited by fibroblasts. J Invest Dermatol. 1999;112(6):835–42.

60. Siersbaek R, Rabiee A, Nielsen R, Sidoli S, Traynor S, Loft A, et al. Transcription factor cooperativity in early adipogenic hotspots and super-enhancers. Cell Rep. 2014;7(5):1443–55.

61. Wu Z, Rosen ED, Brun R, Hauser S, Adelmant G, Troy AE, et al. Cross-regulation of C/EBP alpha and PPAR gamma controls the transcriptional pathway of adipogenesis and insulin sensitivity. Mol Cell. 1999;3(2):151–8.

62. Rosen ED, Hsu CH, Wang X, Sakai S, Freeman MW, Gonzalez FJ, et al. C/EBPalpha induces adipogenesis through PPARgamma: a unified pathway. Genes Dev. 2002;16(1):22–6.

63. Park BO, Ahrends R, Teruel MN. Consecutive positive feedback loops create a bistable switch that controls preadipocyte-to-adipocyte conversion. Cell Rep. 2012;2(4):976–90.

64. Shao D, Rangwala SM, Bailey ST, Krakow SL, Reginato MJ, Lazar MA. Interdomain communication regulating ligand binding by PPAR-gamma. Nature. 1998;396(6709):377-80.

65. Choi S, Jung JE, Yang YR, Kim ES, Jang HJ, Kim EK, et al. Novel phosphorylation of PPARgamma ameliorates obesity-induced adipose tissue inflammation and improves insulin sensitivity. Cell Signal. 2015;27(12):2488–95.

66. Yu W, Chen CZ, Peng Y, Li Z, Gao Y, Liang S, et al. KRAS Affects Adipogenic Differentiation by Regulating Autophagy and MAPK Activation in 3T3-L1 and C2C12 Cells. Int J Mol Sci. 2021;22(24):1–18.

67. Hiemer SE, Varelas X. Stem cell regulation by the Hippo pathway. Biochim Biophys Acta. 2013;1830(2):2323–34.

68. Hong X, Nguyen HT, Chen Q, Zhang R, Hagman Z, Voorhoeve PM, et al. Opposing activities of the Ras and Hippo pathways converge on regulation of YAP protein turnover. EMBO J. 2014;33(21):2447–57.

69. El Ouarrat D, Isaac R, Lee YS, Oh DY, Wollam J, Lackey D, et al. TAZ Is a Negative Regulator of PPARgamma Activity in Adipocytes and TAZ Deletion Improves Insulin Sensitivity and Glucose Tolerance. Cell Metab. 2020;31(1):162–73 e5.

70. Tu Z, Karnoub AE. Mesenchymal stem/stromal cells in breast cancer development and management. Semin Cancer Biol. 2022;86(Pt 2):81–92.

71. Horsley V. Adipocyte plasticity in tissue regeneration, repair, and disease. Curr Opin Genet Dev. 2022;76:101968.

